# *Schistosoma mansoni* antigen induced innate immune memory features mitochondrial biogenesis and can be inhibited by ovarian produced hormones

**DOI:** 10.1101/2025.01.14.632838

**Authors:** Juan Marcos Oviedo, Diana Cortes-Selva, Marco Marchetti, Lauren Gordon, Lisa Gibbs, J. Alan Maschek, James Cox, Seth Frietze, Eyal Amiel, Keke C. Fairfax

## Abstract

We have previously identified that *S. mansoni* infection induces a unique form of myeloid training that protects male but not female mice from high fat diet induced disease. Here we demonstrate that ovarian derived hormones account for this sex specific difference. Ovariectomy of females prior to infection permits metabolic reprogramming of the myeloid lineage, with BMDM exhibiting carbon source flexibility for cellular respiration, and mice protected from systemic metabolic disease. The innate training phenotype of infection can be replicated by *in vivo* injection of SEA, and by exposure of bone marrow to SEA in culture prior to macrophage differentiation (Day 0). This protective phenotype is linked to increased chromatin accessibility of lipid and mitochondrial pathways in BMDM including Nrf1 and Tfam, as well as mitochondrial biogenesis. This work provides evidence that *S. mansoni* antigens induce a unique form of innate training inhibited by ovarian-derived hormones in females.

## Introduction

Cardiovascular disease (CVD) is the leading worldwide cause of mortality (Hinton et al., 2018; Roth et al., 2017). 65% of adults in the United States are diagnosed with diabetes have elevated LDL cholesterol levels or take cholesterol lowering medications, and death rates from atherosclerotic CVD are ∼1.7 times higher in this population as compared to non-diabetic adults (Emerging Risk Factors et al., 2010). It is well established that in the diabetic population, obesity, and dyslipidemia are risk factors underlying these increases in mortality, while hyperglycemia is an independent risk factor (Marks and Raskin, 2000; Wong et al., 2016).

Underlying conditions such as diabetes and atherosclerosis contribute to the burden of CVD in both females and males. While the incidence of CVD is markedly higher in men than in age-matched women (Opotowsky et al., 2007; Tan et al., 2010), the risk of developing CVD while diabetic is much greater in women than men (Humphries et al., 2017; Peters et al., 2014). In non-diabetic patients, females exhibit increased insulin sensitivity in comparison to males, as well as reduced prevalence of dysglycemia and enhanced muscle glucose uptake (Cnop et al., 2003; Kim and Reaven, 2013; Moran et al., 2008; Willeit et al., 1997), suggesting sex-dependent modulations in whole body metabolism. Recent studies suggest gut microbiota contributes to sex differences in lipid metabolism (Baars et al., 2018), but the mechanisms underlying sexual dimorphism in metabolic syndrome remain poorly understood.

Previous studies have uncovered an association between a history of helminth infection and reduced prevalence of metabolic disease in humans and rodents ((Doenhoff et al., 2002; Stanley et al., 2009; Wiria et al., 2015). Specifically, infection by Schistosomes reduces cholesterol and atherosclerotic plaques (Doenhoff et al., 2002; Stanley et al., 2009), this effect has been attributed, in part, to an anti-inflammatory phenotype in macrophages (Wolfs et al., 2014) and transcriptional reprogramming of phospholipid and glucose metabolism related genes in hepatic macrophages (Cortes-Selva et al., 2018b). Moreover, it has been postulated that schistosomes have the potential to affect long term glucose metabolism in T cells (Chen et al., 2013).

Accumulating evidence suggests that biological sex affects disease progression; yet the effect of Schistosomiasis on metabolic-protection in females and males it is not well understood, as most studies have been conducted only in males or no sex differentiation has been made during data analysis ((Sanya et al., 2019; Shen et al., 2015; Wolde et al., 2019)).

Schistosomiasis induces Th2 polarization and alternative activation of macrophages, essential for host survival (Barron and Wynn, 2011; Fairfax et al., 2013; Herbert et al., 2004). IL-4 induced alternative activation of macrophages relies on oxidative phosphorylation (OXPHOS) and fatty acid oxidation for energy production and is dependent on cell intrinsic lysosomal lipolysis

(Huang et al., 2014; Vats et al., 2006). Macrophage metabolism follows a dysmorphic pattern, as sex-related differences affect the processes involved in cholesterol and lipid metabolism in macrophages as well as inflammatory cytokine production in adipose tissue (Griffin et al., 2016; Ng et al., 2001). Moreover, in rats, phagocytes from females had increased ROS generation than males (Rudyk et al., 2018). Our lab recently published that *S. mansoni* infection of male mice induces a form of innate immune memory that modulates myeloid lipid and mitochondrial metabolism, while also protecting from all aspects of high fat diet induced metabolic disease.

Neither this whole-body protection, nor modulation of mitochondrial metabolism occurs in schistosome infected females. Such differences have often been attributed to the role of sex hormones in gene expression and immune cell function (Rubinow, 2018; Taneja, 2018; Winn et al., 2019), but a clear understanding of the effects of sex on the regulation of macrophage metabolism, as well as how sex modulates the effects of Schistosomiasis in the protection from metabolic disease is lacking.

In the present study, we sought to determine the genetic and hormonal mechanisms of sex-dependent schistosome driven innate immune memory. We found profound biological sex driven differences in the transcriptional control of glucose and lipid metabolism, independent of infection status along with differential regulation of key genes in both glycolysis and mitochondrial metabolism by infection in males and females. Surprisingly, we found that while testes produced hormones are not necessary for infection induced protection from metabolic disease in males, elimination of ovarian derived hormones in females allows for schistosome infection to both protect from systemic metabolic disease and induce myeloid metabolic plasticity and innate immune memory. We established that schistosome egg antigens are sufficient to induce protective innate immune training and developed an *in vitro* method for inducing metabolically protective innate immune training that enabled the finding that bone marrow from intact females can be trained *in vitro* without hormones with SEA the same way tht male bone marrow can be . Overall, these data present the first evidence that ovarian produced hormones block metabolically protective schistosome induced innate immune memory and provide a more complete understanding of how schistosome induced innate training may confer metabolic protection at the cellular level.

## Results

### *S. mansoni* infection modulates the myeloid transcriptome in a sex-specific manner

We previously documented significant sex dependent shifts in functional metabolism in BMDM from male and female *S. mansoni* infected mice(Cortes-Selva et al., 2021), we sought to determine if differential transcriptional modulation underlies these shifts. In order to investigate the genes and respective pathways that were associated with specific conditions and the ones that were differentially regulated by sex we performed mRNAseq on unstimulated BMDM derived from male and female ApoE^-/-^ mice at 10-weeks post *S. mansoni* or mock infection. Comparing uninfected males to uninfected females we identified 3926 genes with adj p<0.05 suggesting that there are significant differences in the myeloid compartment of males and females under HFD induced metabolic conditions. Surprisingly, multiple metabolic genes that we identified in our previous publication (Cortes-Selva et al., 2021) as upregulated in males by *S. mansoni* infection are actually downregulated during metabolic disease in uninfected males compared to uninfected females, including *Hk3* (hexokinase 3) , *Mgll* (monoglyceride lipase), *Ptges* (prostaglandin E synthase), and *Slc1a3* (solute carrier family 1 (high affinity glutamate transporter), member 3) (Figure 1A). Specifically comparing BMDM from infected males versus infected females we find that many of these genes that are more highly expressed in females than males during metabolic disease in the absence of infection, are flipped in infected animals, such that *Fabp4* (fatty acid binding protein 4), *Mgll*, *Hk3*, and *Slc1a3* are upregulated in infected males as compared to infected females. Female ApoE^-/-^ mice of reproductive age are similar to female humans in that they do not get as obese or insulin resistant on HFD as male ApoE^-/-^.

**Figure 1.**
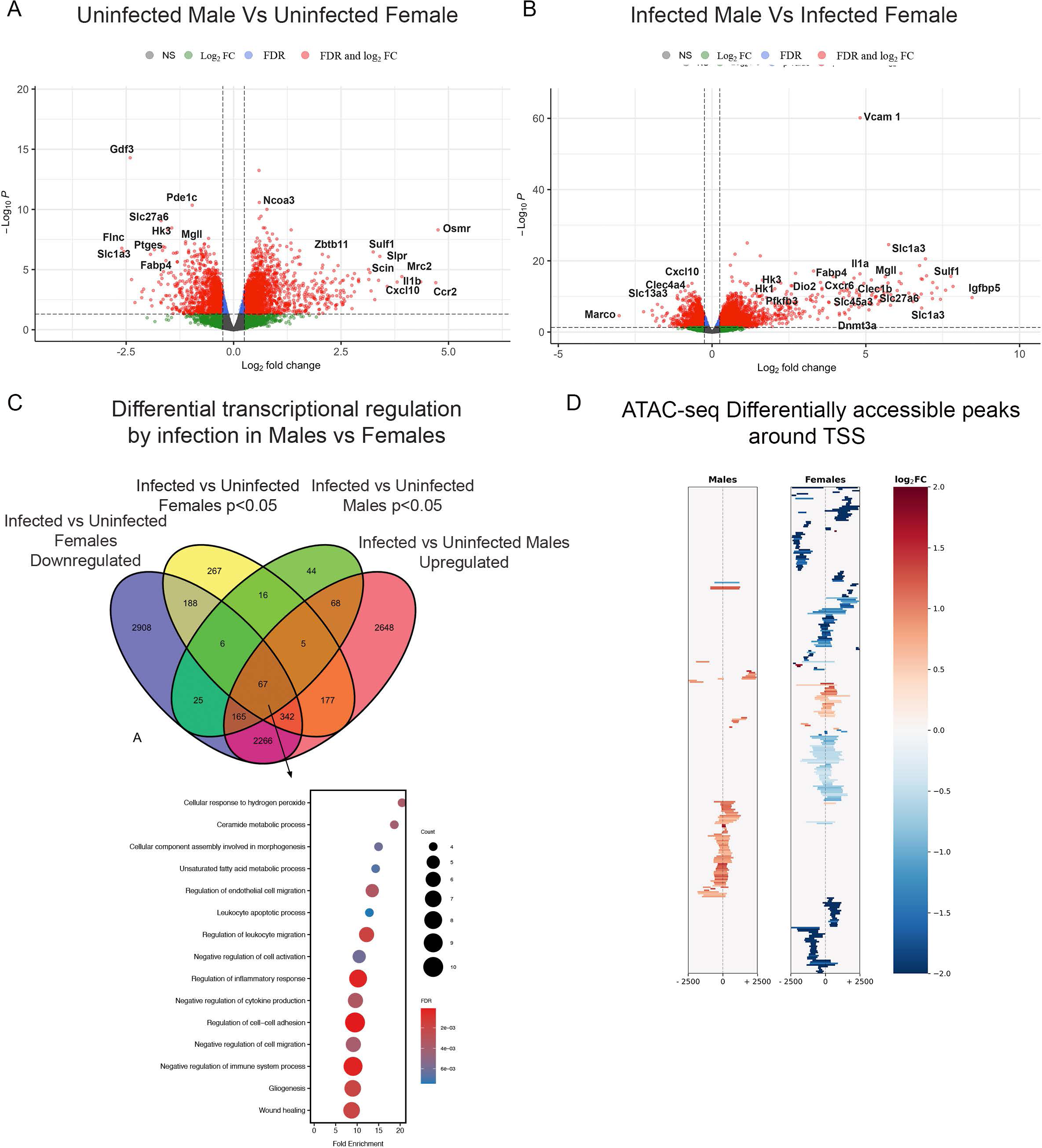
*S. mansoni* infection modulates the myeloid transcriptome in a sex-specific manner. ApoE^-/-^ mice were fed HFD diet for 10-days before infection/mock infection with *S. mansoni*. Bone marrow was harvested at 10-weeks post infection, and BMDM were differentiated with M-CSF. RNA-seq and ATAC-seq was performed on cells from separate cohorts of mice. A) Volcano plot of Genes upregulated in control uninfected ApoE^-/-^ males on HFD versus control uninfected females. B) Volcano plot of Genes upregulated by infection in ApoE^-/-^ males on HFD versus infected females. C) Overlap of genes significantly upregulated by *S. mansoni* infection in males and those downregulated by infection in females (adjusted p-value <0.05) along with pathway analysis of the 67 differentially regulated genes. The gene list is in supplementaryTable .1 D) Heatmap of peaks that are significantly altered by infection near transcription start sites from the ATAC-seq data of each sex. RNA Sequencing data are from one experiment with 4-5 mice per group, ATAC-seq data are from one experiment of 3-4 mice per group.

These data suggest that the regulatory response to infection in the myeloid lineage in males and females is opposite, with males upregulating genes that may be involved in reducing the severity of disease in females, while infection in females downregulates those genes. Critically, our previous work (Cortes-Selva et al., 2021) identified that Mgll activity was needed for increased mitochondrial respiration in BMDM from infected males, as inhibition eliminated the increases in both basal and maximal respiration. We identified different subsets of genes that were preferentially upregulated in males (p<0.05, Supplementary Figure 1A). Among 1444 genes upregulated in males regardless of infection status with a p value <0.05, we identified histone and chromatin remodeling pathways as significantly regulated, suggesting a differential potential for epigenetic regulation in males versus females. A large subset of the 1444 genes (238 genes) were associated to metabolic functions. Of these 238 genes involved in metabolism we identified hexokinase 1 (*Hk1*), citrate synthase (*Cs*), apolipoprotein A2 (*Apoa2*), aldehyde dehydrogenase 3 family member A2 (*Aldh3a2*), lipoyltransferase (*Lipt1*), solute carrier family 19 member 1 (Slc19a1), LDL receptor related protein 1 (*Lrp1*), many of these are involved in lipoprotein and cholesterol metabolism. These data suggest that myeloid metabolism is differentially regulated by sex, at least in the context of HFD induced inflammatory environment. In addition, we found 216 genes involved in immunity. Among genes with immune function, we identified interleukin 10 (*Il-10*), Toll like receptor 5 (*Tlr5*), NLR family pyrin domain containing 3 (*Nlrp3*), inducible T cell costimulatory (*Icos*), which have diverse pro and anti-inflammatory function in the immune system. Next, we surveyed the genes that are differentially regulated in males and females following *S. mansoni* infection. We found 67 genes involved in metabolism (fatty acid and ceramide), hemostasis, the adaptive immune system, cellular migration and activation.

Following the genes with known function, a large fraction of differentially regulated genes have documented roles in metabolism. Among these, we identified type II iodothyronine deiodinase (*Dio2*), which has implicated in the regulation of diet induced obesity (Kurylowicz et al., 2015; Vernia et al., 2013). Moreover, we found hexokinase 3 (*HkIII*), fatty acid binding protein 4 (*Fabp4*), sphingomyelin synthase 2 (sgms2), solute carrier family 6 member 8 (*Slc6a8*), and *Pfkb3* (6-phosphofructo-2-kinase/fructose-2,6-biphosphatase 3) were all upregulated in male and downregulated in female BMDM following *S. mansoni* infection (Figure 1C). PFKFB3 protects against diet-induced adipose tissue inflammatory responses and systemic insulin resistance in mice (Huo et al., 2012), and increased expression of *PFKFB3* is essential for suppressing adipocyte proinflammatory response by metformin (Qi et al., 2017). Importantly, lower PFKFB3 expression in subcutaneous adipose tissue has been associated with insulin resistance in women (Arner et al., 2016). HKIII is not as well studied as HKII, but hexokinase two and three are thought to have some overlapping metabolic functions (Wilson, 2003). HKII overexpression has been shown to increase ATP levels and preserve mitochondrial membrane potential, while also being associated with higher levels of transcription factors that regulate mitochondrial biogenesis, and greater total mitochondrial DNA content (Wyatt et al., 2010). We subsequently performed ATAC-seq analysis on macrophages differentiated from males and females as above. The overall read quality of the uninfected and infected groups was similar across males and females (Supplementary Figure 1B). Focusing on differentially accessible regions that are located around transcription start sites (TSSs), infection in males increases overall accessibility, while infection in females decreases accessibility (Figure 1D), supporting the hypothesis that females, in contrast to males, have an inherently different epigenetic response to schistosomes that likely underlies the lack of a developing a protective metabolic reprograming in response to schistosome infection.

### *S. mansoni*-increases fatty acid oxidation in male but not female ApoE^-/-^ mice on HFD

To determine whether infection-induced protection from weight gain and insulin intolerance in males but not females was accompanied by sex specific differential macrophage metabolic regulation, we cultured bone marrow cells from 10 week infected or uninfected control male and female mice for 7 days to generate BMDM. The Oxygen consumption rate (OCR) was measured in real time in basal conditions and following the addition of mitochondrial inhibitors in unstimulated BMDM from both infected and uninfected male and female ApoE^-/-^ animals.

Consistent with what we have previously published (Cortes-Selva et al., 2021), BMDM from infected males exhibited increased basal OCR and spare respiratory capacity compared to uninfected male controls (Figure 2A, 2B, 2C). However, basal OCR and spare respiratory capacity of BMDM from infected females remained unaltered in comparison to BMDM from uninfected females (Figure 2A, 2B, 2C). Moreover, side by side OCR analysis in females and males with palmitate as a substrate (glucose limiting conditions) showed that BMDM from infected male, but not from infected females had an increased ability to oxidize exogenous palmitate (Figure 2D). In addition, BMDM from infected males but not females had significantly increased palmitate basal OCR and palmitate spare respiratory capacity, suggesting exogenous free fatty usage as a carbon source for OXPHOS (Figure 2E, 2F). Similar to the male only data, we observed no differences in macrophage bulk neutral lipid content in either group, suggesting that global lipolysis may not underlie OCR and spare respiratory capacity in our model (Figure 2G). Additionally, analysis of mitochondrial mass via mitotracker in females and males showed BMDM from male infected mice, but not females have a significantly higher Mitotracker MFI. These data suggest that *S. mansoni* infection induces mitochondrial biogenesis in males, but not females. Again, suggesting differential regulation of macrophage metabolism based on biological sex.

**Figure 2.**
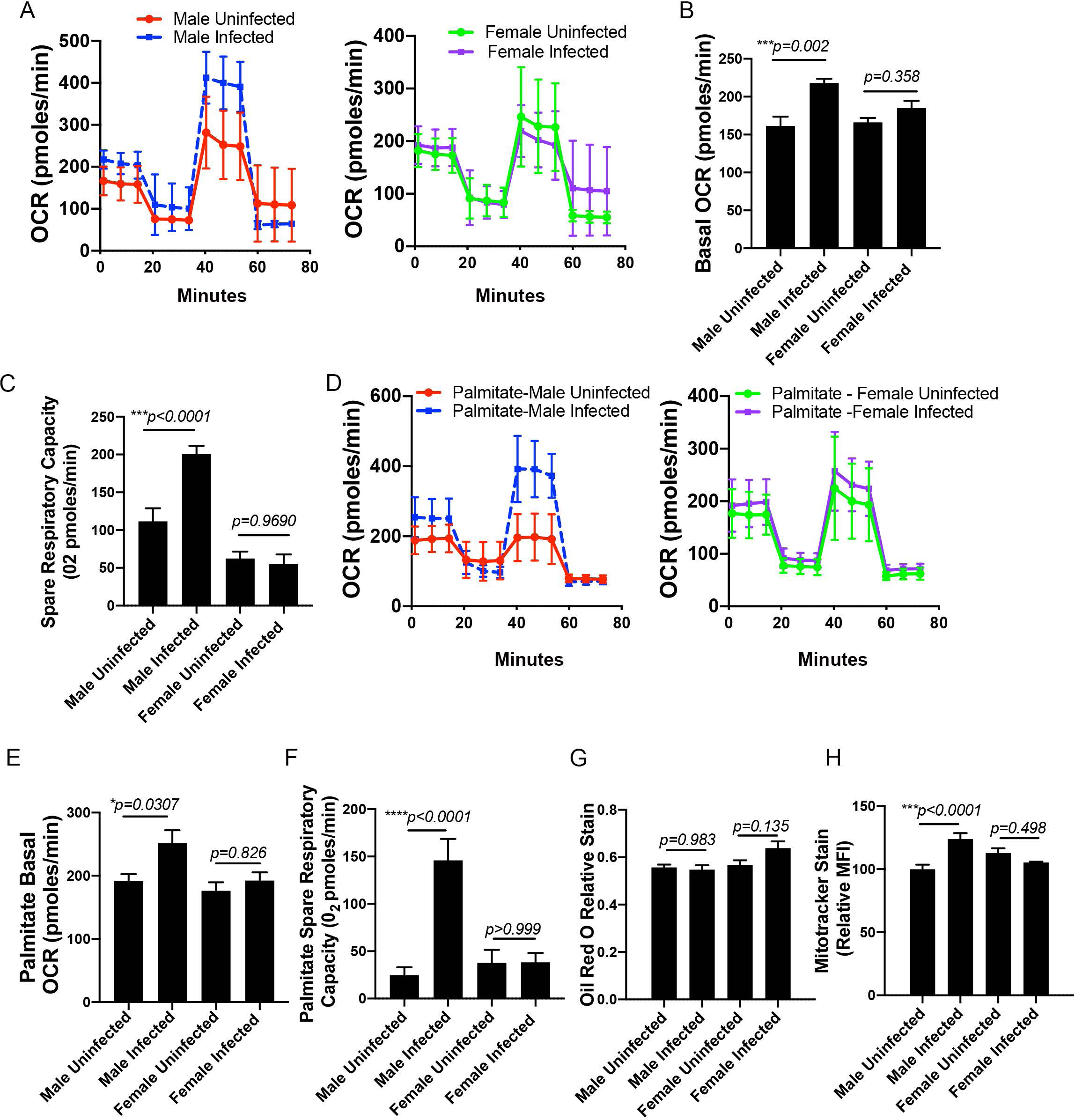
*S. mansoni* increases fatty acid oxidation in male but not female ApoE^-/-^ mice on HFD. Male and female ApoE^-/-^ mice were fed HFD for 10 days before infection with *S. mansoni*. At 10-weeks post infection mice were sacrificed and Mϕ were differentiated from bone marrow with M-CSF. A) SeaHorse assay results for OCR of BMDM from infected and uninfected ApoE^-/-^ males and females in complete media in basal conditions and in response to mitochondrial inhibitors. (B) Quantification (in picomoles/minute) of the basal oxygen consumption of BMDM. (C) Quantification of the spare respiratory capacity of BMDM from A. D) SeaHorse assay results for OCR of BMDM from infected and uninfected ApoE^-/-^ males and females in media where palmitate is the carbon source in basal conditions and in response to mitochondrial inhibitors. E) Quantification (in picomoles/minute) of the basal oxygen consumption of BMDM from D. F) Quantification of the spare respiratory capacity of BMDM from D. G) Oil Red O relative staining in BMDM. H) MitoTracker Red Deep Stain measured by flow cytometry in BMDM.

### *S. mansoni* infection differentially alters the cellular lipid profile in female and male mice

To understand the sex-specific modulations induced by Schistosomiasis in macrophage lipid metabolism, we isolated cellular lipids from unstimulated BMDM and conducted untargeted lipidomics on cells from both male and female mice. Using partial least squares discriminant analysis (PLS-DA), we discerned distinct lipid profiles associated with *S. mansoni* infection in a sex-specific manner (Figure 3A). PLS-DA is relevant as it enhances group separation by maximizing variance explained by the lipidomic data, highlighting significant differences between infected and uninfected samples. Further analysis revealed that total cholesteryl esters (CE), identified as a variable importance in projection (VIP) species in males in our previous publication (Cortes-Selva et al., 2021) decreased in cells from infected male mice but increased in BMDM from infected female mice (Figure 3B). This indicates profound sex-specific metabolic reprogramming in response to infection. Infection in females, similar to males, induces a unique lipid signature (Figure 3C). Notably, plasmanyl-phosphatidylethanolamine (plasmanyl-PE), plasmanyl-phosphatidylcholine (plasmanyl-PC), bis(monoacylglycerol)phosphate (BMP), and CE species were the main drivers of the altered lipid profile in infected females (Figure 3D). Interestingly, BMP abundance remained unchanged by infection in males but increased significantly in females (Figure 3E), suggesting differential regulation of lipid and cholesterol metabolism pathways based on biological sex. Consistent with our previous findings, infection in males increased cellular free fatty acids, while levels remained unchanged in females. These data support the hypothesis that *S. mansoni* infection differentially modulates cellular metabolism in BMDM, with distinct metabolic adaptations in males and females.

**Figure 3.**
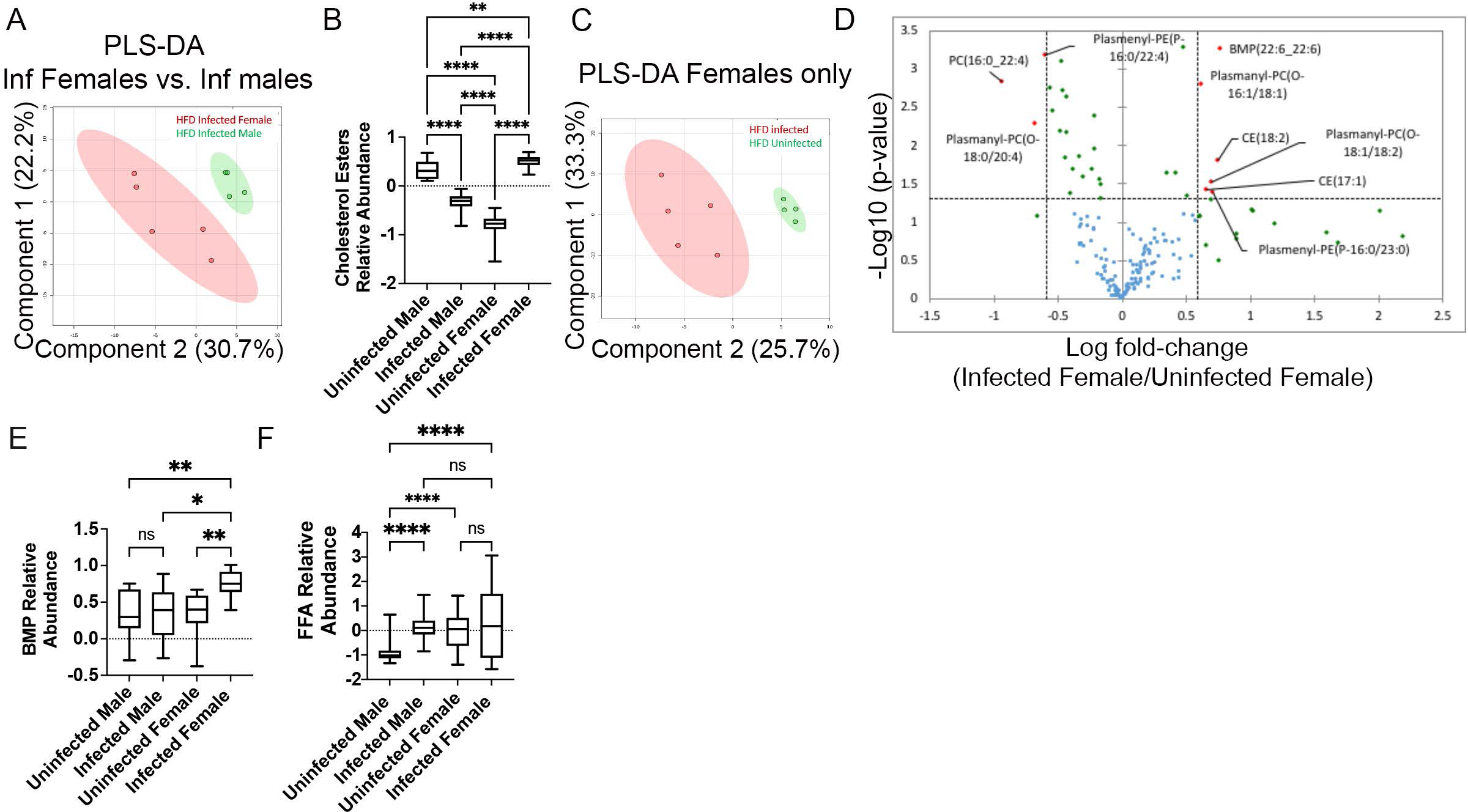
*S. mansoni* infection differentially alters the cellular lipid profile in female and male mice. BM was harvested as above and Mϕs were differentiated with M-CSF in a 7-day culture. Total cellular lipids were extracted and analyzed via LC-MS based lipidomic analysis. A) PLS-DA derived score from Female and Male HFD Infected BMDM. B) Box whisker plot of normalized AUC of cholesterol ester species, between female infected and uninfected ApoE^-/-^ mice on HFD. C) PLS-DA derived score from HFD infected and HFD uninfected female BMDM. D) Volcano plot identifying VIP lipids in infected females. E,F) Box whiskers plots (10–90 percentile) of relative abundance of normalized BMPs and all species of free fatty acids, data are normalized to sum and treated with pareto scaling, each dot is a single species. Data are representative of 3 experiments with 4–6 mice per group in each experiment.

### Elimination of Ovarian hormones allows female to undergo schistosome induced innate training

Considering the strong sex differences in both myeloid transcriptome and metabolic capacity we asked if testes derived hormones are required for protective innate immune training in male, or if ovarian derived hormones in females block schistosome induced innate training. Female ApoE^-/-^ were ovariectomized and males were castrated at 5-6 weeks of age, followed by feeding of high fat diet and infection/mock infection with *S. mansoni*. Ovariectomized uninfected control mice had significantly higher glucose area under the curve and body weight than intact uninfected controls, phenocopying what often occurs after menopause or hysterectomy (Laughlin-Tommaso et al., 2018; Stachowiak et al., 2015). Surprisingly, infected ovariectomized mice had significantly lower glucose AUC than ovariectomized uninfected, intact infected and intact uninfected control females, as well as lower body weight than uninfected ovariectomized mice (Figure 4A). Castration reduces bodyweight somewhat, likely due to the known effects of testosterone on muscle mass (Antonio et al., 1999) but does not eliminate infection induced reductions in glucose area under the curve that reflects better glucose tolerance (Figure 4B). To determine if ovariectomy allowed the same reprogramming of myeloid metabolism as seen in males, we assayed metabolic fuel source flexibility of individual carbon sources (Figure 4C).

**Figure 4.**
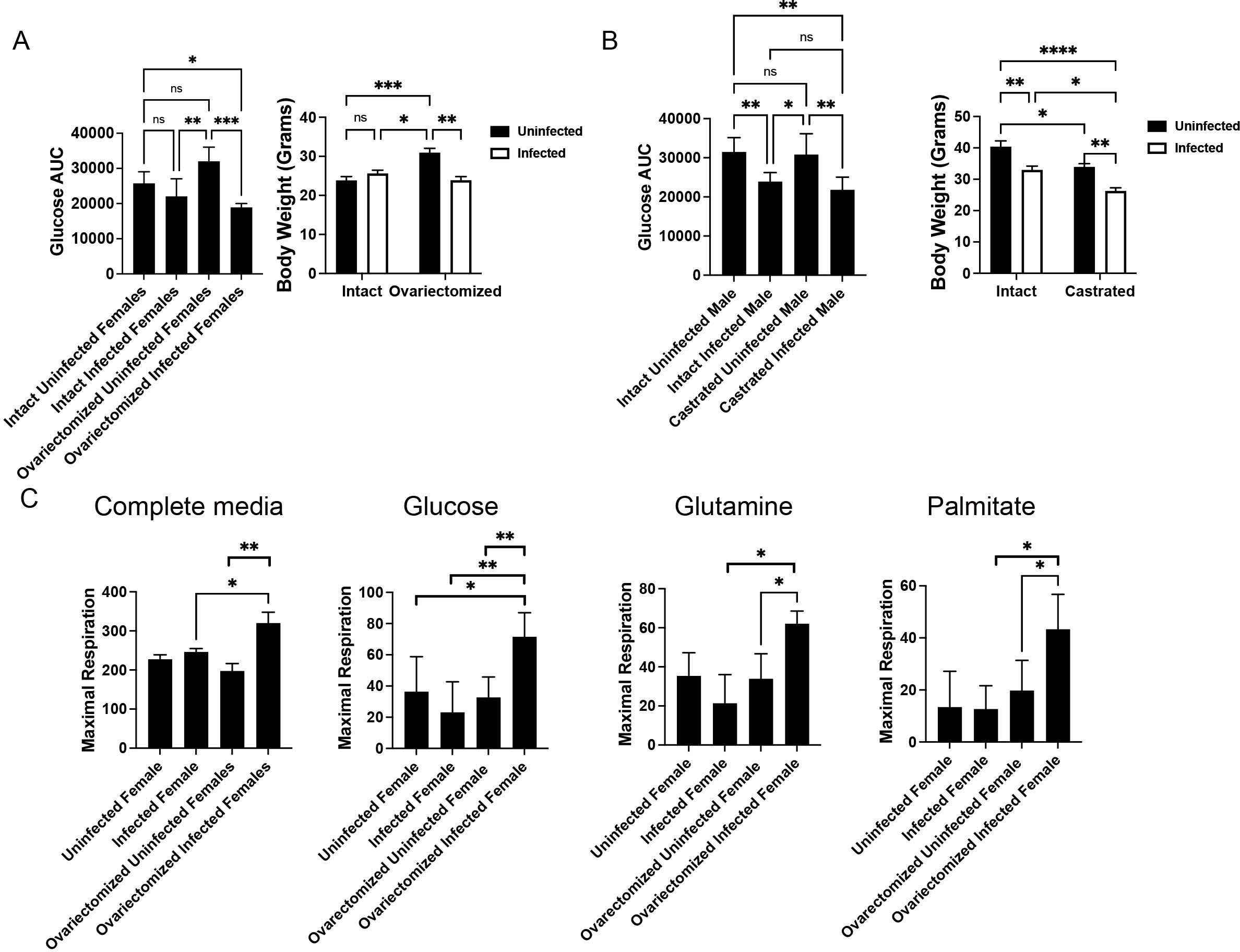
Elimination of Ovarian hormones allows female to undergo schistosome induced innate training. Male and female ApoE^-/-^ mice were gonadectomized at 5.5 weeks of age and then fed HFD for 10 days before infection with *S. mansoni* controls are left intact. At 10-weeks post infection body weight and glucose tolerance were measured prior to sacrifice (A,B). C) Mϕ were differentiated from bone marrow from females in the indicated groups with M-CSF. Maximal respiration for individual carbon sources (Glucose, Glutamine, and Palmitate) measured at steady state in single carbon substrate media and specific inhibitors for each pathway. Data are representative of 3 experiments with 5-6 mice per group in each experiment. Statistical significance calculated using ANOVA.

For the single-carbon substrate extracellular flux assays, data is calculated as the difference between total respiration, and respiration when the specific pathways of interest are inhibited (2-deoxyglucose for glucose, 6-diazo-5-oxo-L-norleucine for glutamine, and etomoxir for palmitate) which reflects a fraction of the total respiration levels observed from BMDM in complete media from ovariectomized infected females. The data demonstrate ovariectomy prior to schistosome infection leads to a significantly greater ability to use all three substrates (glucose, palmitate and glutamine) for cellular respiration, indicating metabolic plasticity similar to what we have published is induced by innate training in males (Cortes-Selva et al., 2021).

### Egg antigen injection is sufficient to induce trained myeloid immunity and protection from metabolic disease in males

Schistosome infection induces dramatic changes to both the systemic cytokine environment and the adaptive immune compartment along with tissue damage that could alter both systemic metabolism and the bone marrow microenvironment. To validate that schistosome egg antigens (SEA) alone can recapitulate the infection induced metabolic protection we have seen in males, we placed male ApoE^-/-^ mice on HFD for 4.5 weeks (to mimic the pre-egg laying period of our infection model) and then began bi-weekly injections of SEA or PBS as a vehicle control. Over the course of 5 weeks of injection, SEA slowed the HFD induced weight gain and after 5 weeks on SEA, treated mice had a 40% reduction in body fat and a 20% increase in lean mass (as measured by NMR, Figure 5A) as well as a significantly lower glucose AUC (Figure 5B). We then assessed whether the myeloid compartment of these mice had undergone training similar to what we have demonstrated for infection by generating BMDM with MCSF in a 7-day culture. BMDM from SEA injected ApoE^-/-^ males have higher OCR area under the curve (Figure 5C) and maximal respiration (Figure 5 D) with palmitate as a carbon source than BMDM generated from control PBS injected mice. These data demonstrate that schistosome egg antigens alone can both modulate whole-body metabolism and induce a unique training of the myeloid linage that features an increase in fatty acid oxidation.

**Figure 5.**
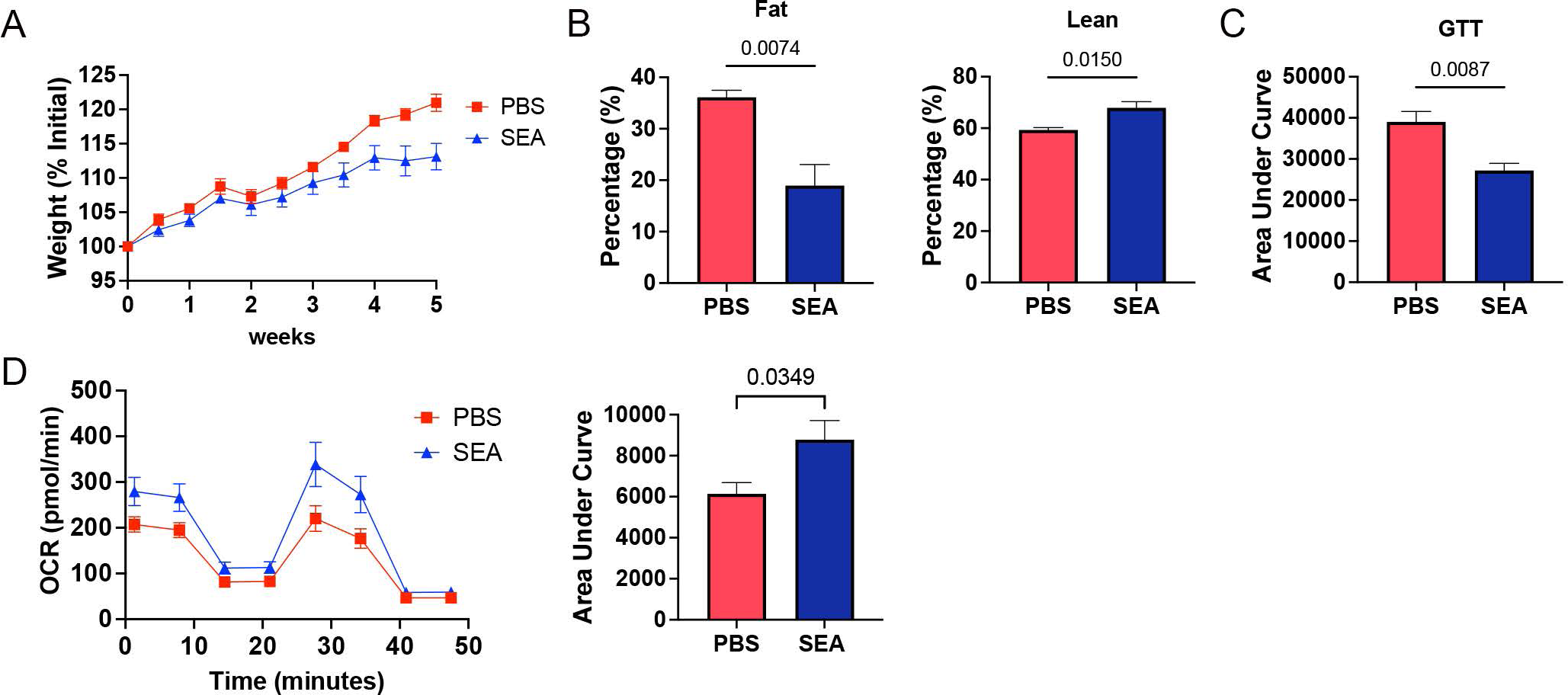
Egg antigen injection is sufficient to induce trained myeloid immunity and protection from metabolic disease in males. Male ApoE^-/-^ mice were fed HFD for 4 weeks before bi-weekly injections of SEA for 5 weeks. Body weight was measured biweekly at the start of SEA administration. B) At 9 weeks post HFD, 5 weeks post SEA-body composition was measured using NMR and a glucose tolerance test was performed. C) SeaHorse assay results for OCR of BMDM from PBS and SEA injected ApoE^-/-^ males in media where palmitate is the carbon source in basal conditions and in response to mitochondrial inhibitors. Data are representative of 2 experiments with 4-5 mice per group in each experiment. Statistical significance calculated using t-tests.

### *In vitro* training of bone marrow progenitors from males and females generates macrophages with metabolic profiles like *in vivo* infection trained cells

Whole body hormonal addback experiments are challenging to use for experiments focused on cellular mechanisms, so we sought to develop a method for inducing schistosome trained immunity *in vitro*. Culture of bone marrow progenitors with SEA prior to macrophage differentiation with MCSF (Day 0) leads to an increase in both basal and maximal respiration that can be further increased in fully differentiated mature macrophages (day 7 of a 7-day culture) with a 24-hour stimulation (SEA d0+24hrs) (Figure 6A, B). Our data in females strongly suggested that ovarian derived hormones may directly block schistosome induced metabolically protective innate training, but it is unclear if this is a chronic or acute effect of the hormones. So we asked if female bone marrow myeloid progenitors could be trained by SEA in culture conditions without hormones. Similar to the male data, female bone marrow trained with SEA *in vitro* prior to macrophage differentiation has a significantly basal (Figure 6 C) and maximal (Figure 6 D) OCR as compared to control untrained cells.

**Figure 6.**
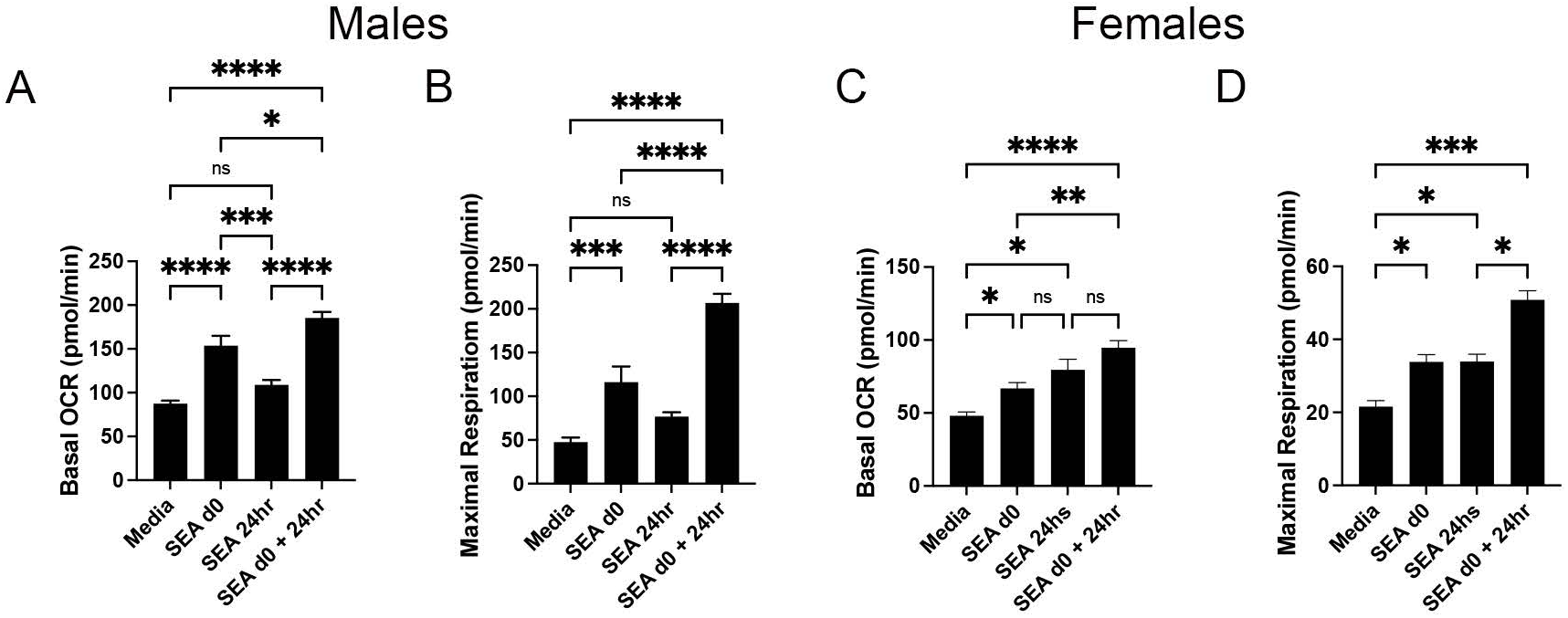
*In vitro* training of bone marrow progenitors from males and females generates macrophages with metabolic profiles similar to infection trained cells. Male and female ApoE^-/-^ mice were fed HFD for 6 weeks before bone marrow was harvested. Bone marrow was exposed to SEA (SEA d0) or just media (No SEA) prior to differentiation of macrophages with M-CSF. At day 7 of differentiation, cells from SEA d0 were split and stimulated with either media or SEA for 24hr (d0+SEA 24hs). SeaHorse assays were performed in complete media. A, C) Quantification (in picomoles/minute) of the basal oxygen consumption of BMDM. B,D) Quantification (in picomoles/minute) of the maximal oxygen consumption of BMDM. Statistical significance calculated using ANOVA data are representative of at least 3 biologically independent experiments.

### SEA innate immune training increases chromatin accessibility of mitochondrial replication and metabolic pathways in male BMDM

Since we can replicate the most salient features of schistosome trained immunity with SEA exposure while removing the influence of systemic inflammation, we performed ATAC-seq on *in vitro* SEA trained BMDM. SEA training prior to macrophage differentiation significantly alters the chromatin accessibility landscape, with lipid metabolic processes being one of the top more accessible pathways (Figure 7A). We identified *Tfam* and *Ffar2* as two genes that have increased chromatin accessibility within the lipid pathways. QPCR analysis showed both genes have increased expression in SEA trained versus control BMDM (Figure 7B). *Ffar2* was also identified in our transcriptomic analysis of BMDM from infected males (Figure 2 and (Cortes-Selva et al., 2021). Ffar2 recognizes short chain fatty acids such as butyrate and has previously been shown to decrease histone deacetylation levels fitting with our data that demonstrates that schistosome trained immunity induces a unique epigenetic profile. Tfam is a nuclear expressed mitochondrial DNA binding protein that has been shown to regulate mitochondrial membrane potential, metabolism (glucose and oxidative phosphorylation, (Koh et al., 2021)) as well as mitochondrial biogenesis in multiple cell types. Using TOBIAS (Bentsen et al., 2020), we performed digital footprinting analysis on ATAC-seq data comparing SEA- and vehicle control trained -macrophages to identify transcription factors with altered accessibility. This analysis (Figure 7C) revealed significant enrichment of TFs involved in mitochondrial biogenesis, metabolic regulation, and immune signaling in SEA-treated cells. Notably, *Nrf1*, a regulator of *Tfam* and a key driver of mitochondrial biogenesis (Zhao et al., 2023), exhibited increased accessibility in SEA-treated macrophages, supporting its role in the metabolic reprogramming observed in schistosome-trained immunity. Additional TFs with increased accessibility included *MLX* and *MLXIPL* (Mejhert et al., 2020), which are involved in lipid and carbohydrate metabolism, as well as members of the E2F family, such as *E2F3* and *E2F4*, known to regulate cell cycle and mitochondrial function (Benevolenskaya and Frolov, 2015). We then compared the TF footprint of infection in females to that of *in vitro* SEA trained male cells. Looking at the most divergent motifs between SEA trained males and infected females, we find the transcription factors we identified in males that regulated mitochondria and cell cycle in Figure 7 C such as *NRF-1* and *E2F4* are uniquely enriched in SEA trained males and not infected females. Motifs that are uniquely enriched for binding in infected females include *Hnf4a*, which can regulate lipid and carbohydrate metabolism (Huck et al., 2021), and *HOXA3* which has been shown to regulated macrophage polarization (Al Sadoun et al., 2016). *MLX* and *MLXIPL* are enriched in both groups compared to their respective controls indicating that they are likely a response to schistosome egg antigens that is unrelated to the metabolic phenotype of enhance mitochondrial respiration. These findings suggest that SEA exposure prior to male macrophage differentiation induces a coordinated epigenetic and transcriptional program to enhance mitochondrial biogenesis and metabolic flexibility.

**Figure 7.**
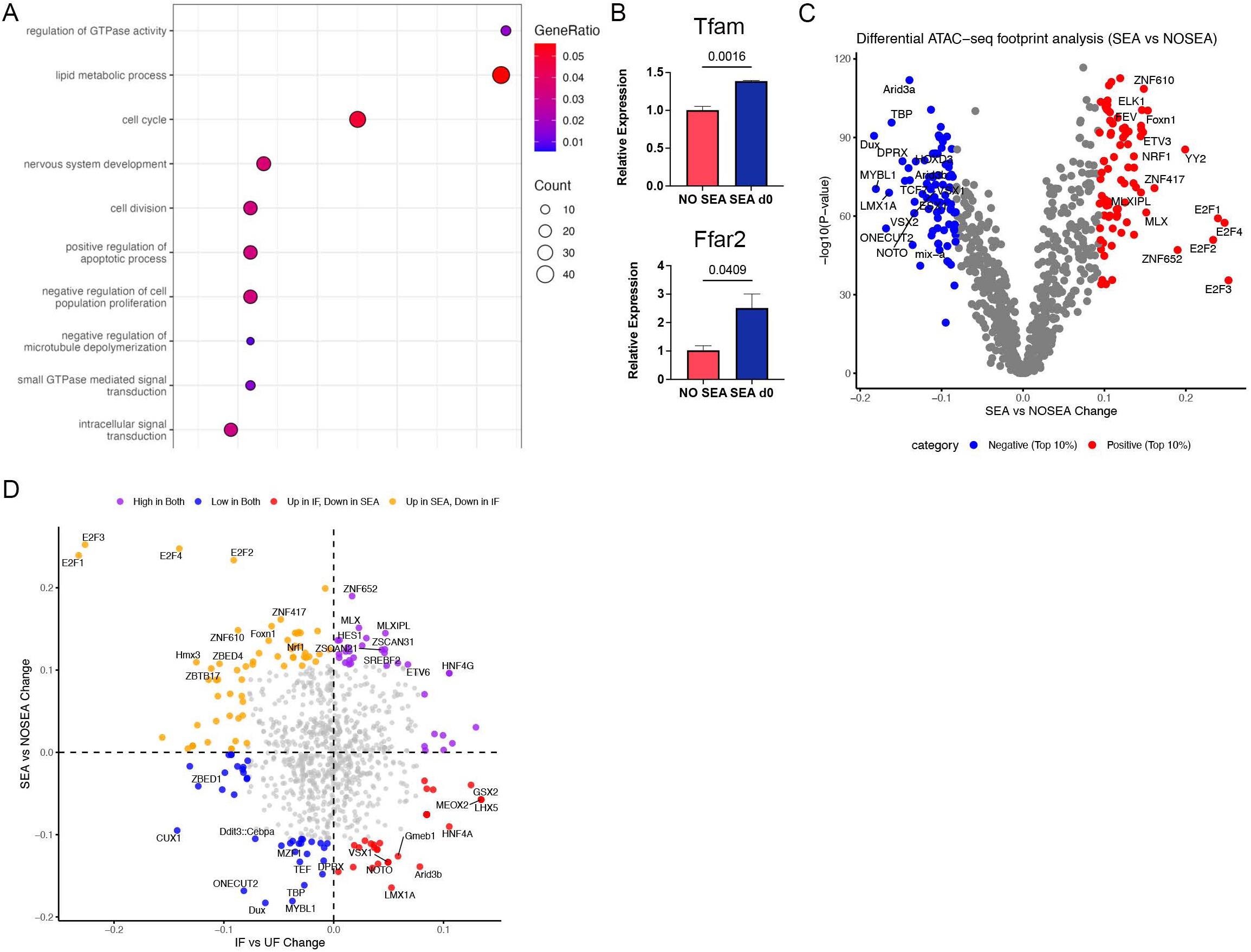
SEA innate immune training increases chromatin accessibility of mitochondrial replication and metabolic pathways. Male ApoE^-/-^ mice were fed HFD for 6-8 weeks before bone marrow was harvested. Bone marrow was trained *in vitro* as described in figure 6 with SEA or PBS control followed BMDM differentiation with MCSF. At day 7 of differentiation nuclei were extracted from cells and ATAC-seq was performed. A) Pathway enrichment analysis of differentially accessible peaks reveals in SEA-trained BMDMs compared to PBS controls. B) Relative transcript abundance of *Tfam* and *Ffar2* quantified by RT-qPCR. Statistical significance was determined using t-tests. C) TOBIAS footprinting analysis of ATAC-seq data identifies TFs with enriched binding activity in SEA-trained BMDMs. D) 4-way scatterplot of the most divergent TF motifs between SEA trained male cells and infected females.

### Schistosome trained immunity increases mitochondrial number in males

Our previous publication that defined schistosome induced innate immune training demonstrated a consistent increase in mitotracker MFI in BMDM from infected males but not females (Figure 2H and (Cortes-Selva et al., 2021)), suggesting that this form of innate immune training may induce mitochondrial biogenesis. To formally evaluate this possibility, we performed electron microscopy (EM) of BMDM generated form the bone marrow of infected and uninfected male and female ApoE^-/-^ mice on HFD. BMDM generated from infected males have a greater number of mitochondria per μm^2^ than control BMDM generated from uninfected males (Figure 8 A, B). Similar to the transcriptional and lipidomic data, schistosome infected females have the opposite phenotype, with fewer mitochondria per μm^2^ than control BMDM from uninfected females.

**Figure 8.**
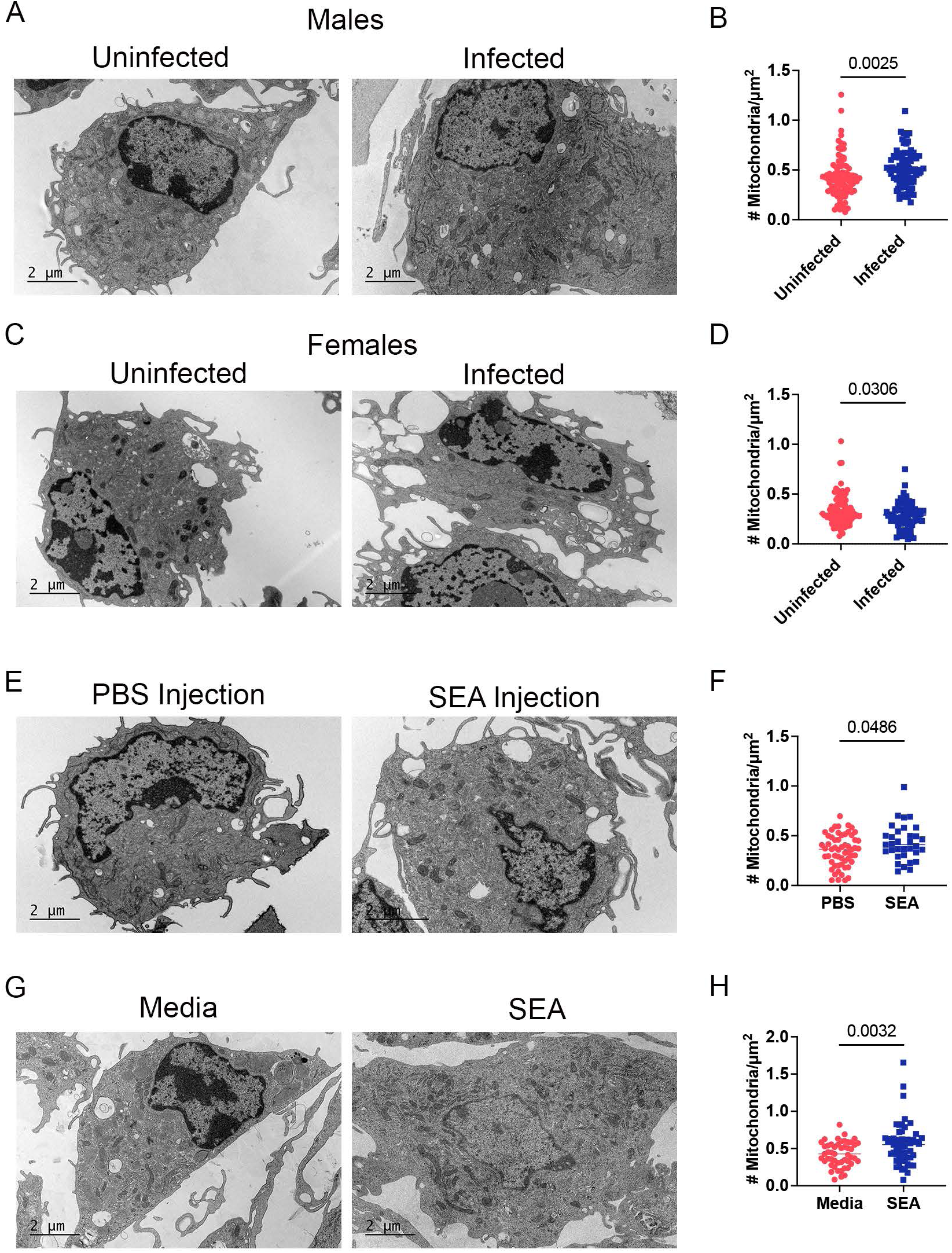
Schistosome trained immunity increases mitochondrial number in males. BMDM were generated as for each condition as described in previous figure. BMDM were fixed with 2.5% glutaraldehyde and 1% paraformaldehyde, processed for embedding in resin, sectioned and then stained with uranyl acetate. Mitochondria were quantified using ImageJ in 1000X images that contained 1-3 macrophages per image. Representative images are 2000x. A, A) BMDM generated from male control and infected ApoE^-/-^ mice on HFD at 10 weeks post infection. C, D) BMDM generated from female control and infected ApoE^-/-^ mice on HFD at 10 weeks post infection E, F) *In vivo* trained SEA Male ApoE^-/-^ mice were fed HFD for 4 weeks before bi-weekly injections of SEA for 5 weeks. G, H) *In vitro* SEA trained ApoE^-/-^ BMDM as described in Fig 6. Statistical significance calculated using t-tests.

Since our data demonstrated that both *in vivo* SEA injection and in vitro SEA training induces a similar metabolic phenotype, we performed EM of BMDM generated from males injected with SEA for 5 weeks and trained *in vitro* in our culture method. Both BMDM generated from SEA injected males (Figure 8E, F) *in vitro* trained BMDM (Figure 8G, H) and have significantly more mitochondria per μm^2^ than control BMDM. Definitively demonstrating that schistosome training of the myeloid lineage involves increases in mitochondrial number and activity.

## Discussion

Helminth infections in general, and Schistosomiasis in specific, have been known to inversely correlate with obesity and glucose intolerance for over a decade, a phenomenon thought to be associated with Type 2 polarization of macrophages and T cells. A cohort in China where Schistosomiasis was recently eradicated, demonstrated that a historical *Schistosoma japonicum* infection is associated with lower prevalence of metabolic syndrome and diabetes (Chen et al., 2013; Shen et al., 2015). There, HbA1c and insulin resistance, as well as triglyceride and LDL levels, were inversely associated with previous *Schistosome* infection, suggesting that not only does active helminth infection modulate metabolic disease, but helminths may induce long-term protection from systemic metabolic disease. Much of the published work of helminth infections and metabolism has focused on males. Our recent study was the first to demonstrate that schistosome infection induces a unique form of innate immune training focused on increased mitochondrial fatty acid oxidation, and that biological sex has a clear effect on the ability of the myeloid lineage to be trained *in vivo* (Cortes-Selva et al., 2021).

Our previous work established that in males, the hallmark of schistosome induced innate training is the generation of a hybrid macrophage that is neither fully polarized to M1 or M2, with decreased production of IL-6 and chronic mediators of inflammation but enhanced transcription of antimicrobial genes and inos. This unique anti-inflammatory phenotype occurs with increased cellular respiration and metabolic plasticity to use multiple carbon sources for fatty acid oxidation respiration. At the cellular lipid level, the hallmark of these cells is dramatically increased free fatty acids with decreased cholesterol esters, TAGs and DAGs. This phenotype appears to be a unique form of innate immune training, as neither BCG nor β-glucan increase fatty acid oxidation, and the hallmark of those forms of trained immunity in increased inflammation, IL-6 and glycolysis (Arts et al., 2016). The long-lived natures of both BCG and β-glucan trained immunity are enforced via epigenetic rewiring of bone marrow progenitors, with BCG and β-glucan inducing enrichment of H3K4me3 at the promoters of IL6 and TNF, and BCG additionally enriching H3K9me3 (Arts et al., 2016). IL-6 production/secretion is enhanced in both BCG and β-glucan innate training neither of which are known to be metabolically protective. Intriguingly, *Dnmt1*, involved in DNA methylation maintenance (Hervouet et al., 2018), is downregulated in males but remains statistically unchanged in females in our transcriptomic analysis following infection. While *Dnmt3a*, essential for establishing DNA methylation patterns during development and implicated in hematopoietic stem cell clonal expansion (Wang et al., 2024), is upregulated in males but downregulated in females (Figure 1). These data support the hypothesis that sex-specific differences in the regulatory epigenetic landscape underlie the protective metabolic reprogramming observed in males but not females.

In our model the most direct comparison for alterations to the chromatin landscape between schistosomes and β-glucan would be the ATACseq data from *in vitro* SEA trained BMDM. A large percentage of the regions with increased accessibility are related to mitochondrial function and cell cycle, which have not been reported for either BCG or β-glucan. Indeed, one of the transcription factors that is enriched from the TOBIAS analysis is Nrf1, which regulates *Tfam* expression and pathways related to mitochondrial respiratory function (Satoh et al., 2013). We validated that *Tfam* transcript is indeed upregulated in these cells (Fig 7). Tfam has been shown to increase hexokinase activity (Koh et al., 2019), so we hypothesize that the combined effects of epigenetically upregulating *Nrf1* and *Tfam* are key components of metabolically protective schistosome induced innate training. Critically, *Nrf-1* and *E2f* family motif enrichment does not occur in infected females (Figure 7 D), which do not have increases in number of mitochondria or in mitochondrial respiration.

We have previously hypothesized that schistosome induced training increases mitochondrial respiration in males via mitochondrial biogenesis, but that hypothesis was based only on flowcytometric quantitation of mitotracker staining. In this current study we performed electron microscopic imaging of BMDM generated from infected and control males and females and found that BMDM from infected males have increased mitochondria per area compared to control uninfected, while females have fewer mitochondria. Additionally, on the 2000x images it appears that BMDM from infected females may have more small lipid bodies, although we have not formally quantified that, future work will focus on these sex specific ultrastructure differences. Since schistosome infection induces systemic inflammation and increase to many cytokines, we utilized *in vitro* and *in vivo* SEA exposure to determine if the increase in mitochondria in male BMDM were directly driven by schistosome antigens. BMDM from both training conditions demonstrate increased mitochondria, with the most dramatic increases in the culture trained BMDM, definitively indicating that SEA training of the myeloid lineage induces mitochondrial biogenesis.

In the current study we report that schistosome infection differentially regulates the transcriptome, chromatin landscape, and mitochondrial metabolism in males and females and that schistosome egg antigens exposure can replicate this unique innate immune training *in vivo* and *in vitro*. Analysis of the BMDM transcriptomes from males and females revealed that over a thousand genes are upregulated in male versus female BMDM regardless of *S. mansoni* infection state. Hundreds of these are annotated to metabolic pathways, suggesting that there are inherent differences in male and female myeloid cellular metabolism. Over five dozen are differentially regulated by infection. Focusing on the genes that are upregulated in BMDM from infected males and downregulated in females, we found more than ten genes with known functions in cellular metabolism. Some of these genes, like *PFKFB3*, have known regulatory elements for progesterone and estrogen, but some of them, like *Fabp4*, have no published mechanism of sex hormone regulation. Hexokinase transcripts are elevated in males and either not changed (*Hk1*) or downregulated (*Hk3*) in females. Estrogen has been documented to regulate hexokinase enzyme activity (but not transcription) in some cell types via activation of the Akt pathway (Brinton, 2008). It is unclear if that occurs during schistosome infection as glycolysis is not significantly increased in infected females. Our previously published data indicates that for males, the upregulation of glycolysis is to generate TCA cycle intermediates and not lactate as an end product (Cortes-Selva et al., 2021), so the regulation induced by infection may be distinct from other models. Our data from ovariectomized and castrated mice demonstrate that the sex difference is most likely driven by an inhibitory effect of ovarian derived hormones, as castration of males does not prevent schistosome induced training, but ovariectomy of females renders them permissive to schistosome induced training. Indeed, our culture-based method of inducing schistosome innate training demonstrates that in culture conditions where physiological levels of female hormones are absent, female bone marrow can generate trained BMDM with increased mitochondrial respiration levels similar to males. Combined with our ATAC-seq data showing that infection increases chromatin accessibility in males and decreases it in females (Fig 1), there is strong support for the hypothesis that female hormones inhibit schistosome induced training at the progenitor level.

At the whole-body level there are known differences in lipid and glucose metabolism between males and females, with females generally having more body fat (higher subcutaneous and lower visceral) as well as lower muscle mass. The regulation of the distribution of body fat has been attributed to estrogen, as deposition shifts from subcutaneous to visceral after menopause (Svendsen et al., 1995) and deletion of estrogen receptor in mice leads to less subcutaneous adiposity (Lapid et al., 2014). Additionally, women tend to have higher non-oxidative free fatty acid clearance via re-esterification than men (Nielsen et al., 2003), suggesting that women tend to store fatty acids from the circulation while men oxidize them. At the myeloid level, males on high fat diet have increased myelopoiesis compared to females, as well as more proinflammatory adipose tissue macrophages (Singer et al., 2014). These differences are hormonally regulated as castration of males in this model reduces adipose monocyte inflammation while ovariectomy of females increased adipose tissue macrophages and bone marrow myeloid colony formation (Varghese et al., 2021). Very few studies have focused on sex differences in myeloid cellular lipid or glycolytic metabolism but comparing the uninfected control male and female lipidomics in figure 3, females have a higher abundance of free fatty acids in addition to multiple PE and PC species than males. While our 2018 publication (Cortes-Selva et al., 2018a) did not include females, the liver macrophage transcriptomic data suggested similar shifts in male myeloid cellular metabolism as we have described in BMDM in both this current paper and our 2021 paper (Cortes-Selva et al., 2021). Taking all these data together, the scientific premise that there is a flip in storage versus usage of lipids and free fatty acids in male and female macrophages that is amplified in the context of schistosome induced innate immune training is well supported. Additional studies are needed to determine if our BMDM data is representative of adipose and other tissue macrophage population during metabolic disease in males versus females and if schistosome induced innate training similarly drives fatty acid oxidation in metabolic tissues.

There are significant differences in both the susceptibility to disease presentation of diabetes and CVD between males and females, with females having lower rates of diabetes but increased risk of cardiovascular complications once diabetic (Humphries et al., 2017; Peters et al., 2014; Yoshida et al., 2023). Few studies have focused on the role of immunometabolism in this dichotomy. Females of reproductive age are generally protected from severe metabolic disease compared to similarly obese males (Li et al., 2022). This protection goes away post-menopause (Heianza et al., 2013), suggesting that ovarian derived hormones are involved. Data from cancer, type 1diabetes, and reproductive studies indicates that progesterone can modulate whole body and tissue/cellular glucose metabolism (Atif et al., 2019; Picard et al., 2002) with sometimes conflicting results (Lee et al., 2020; Lee et al., 2018). Estradiol can both bind directly to fatty acids and induce a form of lipolysis, while also increasing glucose uptake *in vitro*, and hepatic gluconeogenesis *in vivo* in some models (Yan et al., 2019). Estradiol replacement therapy in post-menopausal women lowered blood glucose and increased insulin sensitivity in some studies (Friday et al., 2001), while others found an increasing risk of type 2 diabetes with increasing duration of postmenopausal estrogen use (Zhang et al., 2002). Our data suggests clear biological sex differences driven by ovarian derived hormones in both the ability of schistosomes to protect from the development of HFD induced metabolic disease parameters, and the ability of schistosome antigens to train the myeloid lineage towards a protective phenotype via mitochondrial biogenesis and increased respiration. There are limitations to our current model, as SEA contains hundreds of proteins as well as carbohydrates and lipids. Future work will be focused on determining what components of SEA are necessary and sufficient to induce the metabolically protective innate training described in this study. Our current data also indicate a clear need for further studies in both humans and animal models to specifically probe the relationship between biological sex and myeloid metabolism, and how various steroid hormones regulate both cellular and whole-body metabolism.

## Acknowledgements

KCF and EA conceived of the project; KCF, EA, JEC, JMO, and DCS designed the experiments; KCF, DCS, LG, EA, and JAM, performed the experiments; KCF, DCS, EA, JAM, JEC, and SF analyzed the data; KCF, DCS, and JMO wrote the manuscript. The work was supported by The University of Utah, a Scientist Development Grant from the American Heart Association to KCF (14SDG18230012), 1 R01 AI158710-01 to KCF and EA, an American Heart Association Pre-doctoral Award (18PRE34030086) to DCS, and 1R21AI135385-01A1 to EA. *B. glabrata* snails provided by the NIAID Schistosomiasis Resource Center of the Biomedical Research Institute (Rockville, MD) through NIH-NIAID Contract HHSN272201700014I for distribution through BEI Resources. JEC is supported through U54 DK110858-01, mass spectrometry equipment employed was provided by 1S10OD016232-01, 1S10OD018210-01A1 and 1S10OD021505-01 to JEC.

## LEAD CONTACT AND MATERIALS AVAILABILITY

Further information requests should be directed to and will be fulfilled by the Lead Contact, Keke Fairfax. These studies generated no new reagents.

## EXPERIMENTAL MODEL AND SUBJECT DETAILS

### Ethics statement

This study was carried out in accordance with the recommendations in the Guide for the Care and Use of Laboratory Animals of the National Institutes of Health. The protocols were approved by the Institutional Animal Care and Use Committees of the University of Utah (#18– 09001).

### Parasite and mouse models

Snails infected with *S*. *mansoni* (strain NMRI, NR-21962) were provided by the Schistosome Research Reagent Resource Center for distribution by BEI Resources, NIAID NIH. ApoE^-/-^ (B6.129P2-Apoetm1Unc/J) were purchased from the Jackson Laboratories and bred at the University of Utah. Castration and Ovariectomy or sham-operation were performed by Jax surgical services. 6-8-week-old male mice were housed in pathogen-free conditions and were fed standard rodent chow (2019 rodent chow, Harlan Teklad) until 10–14 days before infection when they were transitioned to a high-fat diet (HFD: 21% milk fat, 0.15% cholesterol: TD 88137 Envigo).

### *S*. *mansoni* infection and glucose tolerance test

ApoE^-/-^ male mice of 6 weeks of age were exposed percutaneously to 75–90 cercariae of *S*. *mansoni* or were mocked infected (as controls). At five- and ten-weeks post-infection mice were fasted for four hours and baseline blood glucose levels were obtained via lateral tail vein nick. Mice were then administered a single intraperitoneal injection of glucose (2mg/g of body weight, ultrapure glucose, Sigma G7528). Blood glucose levels were obtained at 20, 60, and 90-minutes post injection. Individual data points obtained were analyzed by Area Under Curve (AUC).

### SEA injections

Male or female ApoE^-/-^ of 6 weeks were on HFD for 4 weeks. After that, mice were injected biweekly with 50 ug of SEA or PBS as negative control and weight variation measured. After 5 weeks of injections, an IP glucose tolerance test was performed (Cortes-Selva et al., 2021). Body weight and fat were measured using a Bruker Minispec LF50 device (Bruker, Karlsruhe, Germany). Finally, bones were collected for cultures to generate BMDM.

### Mouse macrophage culture

Mouse bone marrow-derived macrophages (BMDM) were generated as follows: bone marrow cells were isolated by centrifugation of bones at >10,000 x g in a microcentrifuge tube for 15 seconds as previously described [<u>76</u>]. Cells were differentiated in M-CSF (20ng/mL, Peprotech, Rocky Hill, NJ) in complete macrophage medium (CMM: RPMI1640, 10% FCS, 2mM L-glutamine and 1 IU/mL Pen-Strep for 6 or 7 days. On the last day, cells were harvested in Cellstripper cell dissociation reagent (Corning) were washed with CMM and prepared for downstream assays. For generation of ex vivo SEA trained macrophages, 5ug/ml of SEA was added to the complete media culture with the bone marrow cells at (d0), for stimulation of mature macrophages, the same amount of SEA was added at d6. When the effect of sexual hormones was tested charcoal stripped Fetal Bovine serum (Gibco, Qualified One shot) and media without phenol red were used.

### Flow cytometry

Staining of BMDM was performed using the following mAb against mouse antigens: F4/80 (BM8, Biolegend), Mitotracker DeepRed (ThermoFisher), and Oil RedO (Thermo Fisher) . Samples were acquired using Attune NxT Focusing Flow Cytometer (Thermo Fisher Scientific) and analyzed using Flowjo X 10.10.0 (FlowJo LLC, Inc.).

### Glycolytic and phospho-oxidative metabolism measurement (seahorse assay)

BMDM from different conditions were resuspended at the same concentration in XF assay media supplemented with 5% FCS and 5mM glucose. The day before the assay, the probe plate was calibrated and incubated at 37 C in a non-CO2 incubator. Resuspended cells were seeded at a concentration of 1.5x10^5^ cells per well and incubated for 20-60 minutes in the Prep Station incubator (37 C non-CO2 incubator). Following initial incubation, XF Running Media (XF assay media with 5% FCS and 10mM Glucose) were dispensed into each well. OCR and ECAR were measured by an XF96 Seahorse Extracellular Flux Analyzer following the manufacturer’s instructions. For the seahorse assay, cells were treated with oligomycin (1uM), FCCP (1.5uM), rotenone (100nM) and antimycin A (1uM). Each condition was performed in 2-3 technical replicates. For determination of palmitate dependent respiration, BSA-conjugated palmitate (BSA: palmitate = 1:6, molar ratio) was prepared according to the Seahorse protocol (Seahorse Bioscience). Briefly, 1 mM sodium palmitate (Sigma Aldrich) was conjugated with 0.17 mM fatty acid free-BSA (Sigma Aldrich) in 150 mM NaCl solution at 37°C for 1h. Palmitate-BSA was stored in glass vials at -20°C until use. Cells were incubated as above in glucose limited XF media per manufacturer instructions.

### RNA Isolation and q-RT-PCR

BMDM were washed with PBS and then lysed with 350 ul of LBP (Takara). Then, Nucleospin RNA plus kit (Takara) was used to extract RNA following manufacturer instructions. Next, cDNA was synthesized from RNA using Superscript IV VILO (ThermoFisher Scientific) for reverse transcription. qPCR was performed using TaqMan Gene expression assays (mgll, slc1a3, beta actin, ThermoFisher) on an Applied Biosystems Stepone Plus Real-Time PCR System.

Beta-Actin assay number Mm02619580_g1, Tfam assay Mm00447485_m1, Ffar2 assay Mm02620654_s1. Relative expression was calculated using the 2-ΔΔCt method (Arocho et al., 2006).

### RNA Sequencing

The sequencing data were first assessed using FastQC (Babraham Bioinformatics) for quality control. Then all sequenced libraries were mapped to the mouse genome (UCSC mm10) using STAR RNA-seq aligner [<u>77</u>] with the following parameter: “—outSAMmapqUnique 60”. The reads distribution across the genome was assessed using bamutils (from ngsutils) [<u>78</u>]. Uniquely mapped sequencing reads were assigned to mm10 refGene genes using featureCounts (from subread) [<u>79</u>] with the following parameters: “-s 2 –p–Q 10”. Quality control of sequencing and mapping results was summarized using MultiQC (Ewels et al., 2016). Genes with read count per million > 0.5 in more than 2 of the samples were kept. The data was normalized using trimmed mean of M values method. Differential expression analysis was performed using edgeR (Robinson et al., 2010). False discovery rate was computed from p-values using the Benjamini-Hochberg procedure. Pathway analysis was performed on log2-transformed data using Bonferroni-corrected *p*-values. The data discussed in this publication have been deposited in NCBI’s Gene Expression Omnibus (Edgar et al., 2002) and are accessible through GEO Series accession number GSE144447 (https://www.ncbi.nlm.nih.gov/geo/query/acc.cgi?acc=GSE144447).

### ATAC-seq

BMDM were treated with lysis buffer with digitonin to extract nuclei. Then, approximately 50000 nuclei were used for tagmentation following manufacturer’s instructions (Diagenode), DNA purified, and libraries were prepared. After QC, libraries were sequenced using novogene service using NovaSeq 6000. Raw sequencing reads were pre-processed by trimming adapters and removing low-quality reads with TrimGalore (0.6.10) prior to alignment to the mouse genome (GRCm39.113) using BWA (0.7.17). After alignment duplicate reads and mitochondrial reads were removed. Chromatin accessibility peaks were called using MACS2 (2.2.9.1) with a q-value cutoff of 0.01. The resulting peaks were processed with BEDTools (2.31.1) to create a consensus set, then annotated to the nearest genes using HOMER (4.11). Peak reads were counted using featureCounts from the Subreads software package (2.0.6), then counts were used for differential accessibility analysis using the R package DESeq2 (1.42.1). For pathway enrichment analysis, genes were first filtered by removing those with discordant associated peaks (*i.e.* genes having both opened and closed peaks). For bias correction and transcription factor footprinting, we applied TOBIAS to the ATAC-seq data (Bentsen et al., 2020). Footprint scores were calculated using TOBIAS ScoreBigwig and differential motif binding was analyzed using TOBIAS BINDetect and JASPAR 2022 motif libraries to estimate TF binding activities. ATAC-seq data will be deposited in GEO, accession number available at publication.

### Untargeted Lipidomics Workflow

#### Sample extraction from serum or cell pellets

Lipids are extracted from serum (50uL) or cell pellets in a combined solution as described in (Matyash et al., 2008). Samples were incubated with 225 µL methanol (MeOH) containing internal standards (IS; Avanti Splash Lipidomix, 10 µL per sample) and 750 µL methyl tert-butyl ether (MTBE). The samples were sonicated for 1 minute, rested on ice for 1 hour with brief vortexing every 15 minutes, followed by the addition of 200 µL double-distilled water (ddH₂O) to induce phase separation. The samples were then vortexed for 20 seconds, rested at room temperature for 10 minutes, and centrifuged at 14,000 g for 10 minutes at 4°C. The upper organic phase was collected and evaporated to dryness under vacuum. Lipid samples were reconstituted in 200 µL isopropanol (IPA) and transferred to an LC/MS vial with insert for analysis. A process blank sample and a pooled quality control sample (10 µL per sample) were also prepared.

#### LC-MS Methods

Lipid extracts were separated on a Waters Acquity UPLC CSH C18 1.7 µm 2.1 x 100 mm column maintained at 65°C, connected to an Agilent HiP 1290 Sampler, Agilent 1290 Infinity pump, and Agilent 6530 Accurate Mass Q-TOF dual ESI mass spectrometer. For positive mode, the source gas temperature was set to 225°C, with a gas flow of 11 L/min and a nebulizer pressure of 50 psig. For negative mode, the source gas temperature was set at 325°C, with a drying gas flow of 12 L/min and a nebulizer pressure of 30 psig. Samples were analyzed in a randomized order in both positive and negative ionization modes in separate experiments, acquiring with the scan range m/z 100-1700. Mobile phase A consisted of acetonitrile:water (60:40 v/v) with 10 mM ammonium formate and 0.1% formic acid, and mobile phase B consisted of isopropanol:acetonitrile:water (90:9:1 v/v) with 10 mM ammonium formate and 0.1% formic acid. The chromatography gradient started at 15% mobile phase B, increased to 30% B over 2.4 minutes, then to 48% from 2.4-3.0 minutes, followed by an increase to 82% B from 3-13.2 minutes, and then to 99% from 13.2-13.8 minutes, where it was held until 15.4 minutes before returning to the initial condition and equilibrating for 4 minutes. The flow rate was 0.4 mL/min throughout, with an injection volume of 5 µL for positive mode and 7 µL for negative mode. Tandem mass spectrometry was conducted using the same LC gradient at collision energies of 20 V and 40 V.

#### Data Analysis and Pretreatment

Pooled quality control (QC) samples and process blanks were injected throughout the sample queue to ensure the reliability of the acquired lipidomic data. Results from LC-MS experiments were collected using Agilent Mass Hunter (MH) Workstation and analyzed using the software packages MH Qual, MH Quant (Agilent Technologies, Inc.), and LipidMatch (Koelmel et al., 2017) to prepare the data set. The data table exported from MH Quant was evaluated using Excel, where initial lipid targets were parsed based on the following criteria: only lipids with a relative standard deviation (RSD) of less than 30% in QC samples were used for data analysis. Additionally, targets identified in blanks or double blanks at significant amounts (area under the curve (AUC) target blank/AUC target QC >30%) were removed from analysis.

##### Electron Microscopy Imaging

BMDMs were washed with PBS and fixed using Fixatives solutions (Electron Microscopy Sciences) overnight, stored at 4°C in fixative solution and sent to the Electron Microscopy Laboratory at University of Utah for further processing. After postfixation with 2% osmium tetroxide and prestaining with uranyl acetate, the ventricular tissue slices were dehydrated in graded ethanol and pure acetone, imbedded with epoxy resin, and sectioned at 70 nm using an ultramicrotome (Leica). Sections were poststained with acetate and lead citrate before imaging with JEM-1400Plus or JEM1200-EX (JEOL) transmission electron microscope with a CCD Gatan camera.

)Samples were then dehydrated using a graded ethanol series and embedded in Embed 812 (Electron Microscopy Sciences) using a graded resin and propylene oxide series. The 1-μ transverse sections were cut with a diamond knife ultramicrotome and imaged using a Jeol 1400-plus transmission electron microscope (Jeol, Tokyo, Japan) with a high-resolution digital camera (AMT, Woburn, MA, USA). Images were captured at 1000 and 2000x for counting mitochondria and for representative images for publication. Mitochondria were counted in ImageJ.

### Statistical Analysis

Statistical analyses of data were performed using one-way ANOVA, a non-parametric Mann-Whitney test, or unpaired Student’s t-test depending on the data distribution. P ≤ 0.05 were considered statistically significant. Analyses and graphing were performed using Prism (GraphPad v10.0) and R-language for statistical computing.

