## Supplemental Figure 1 for "*Schistosoma mansoni* antigen induced innate immune memory features mitochondrial biogenesis and can be inhibited by ovarian produced hormones"

A

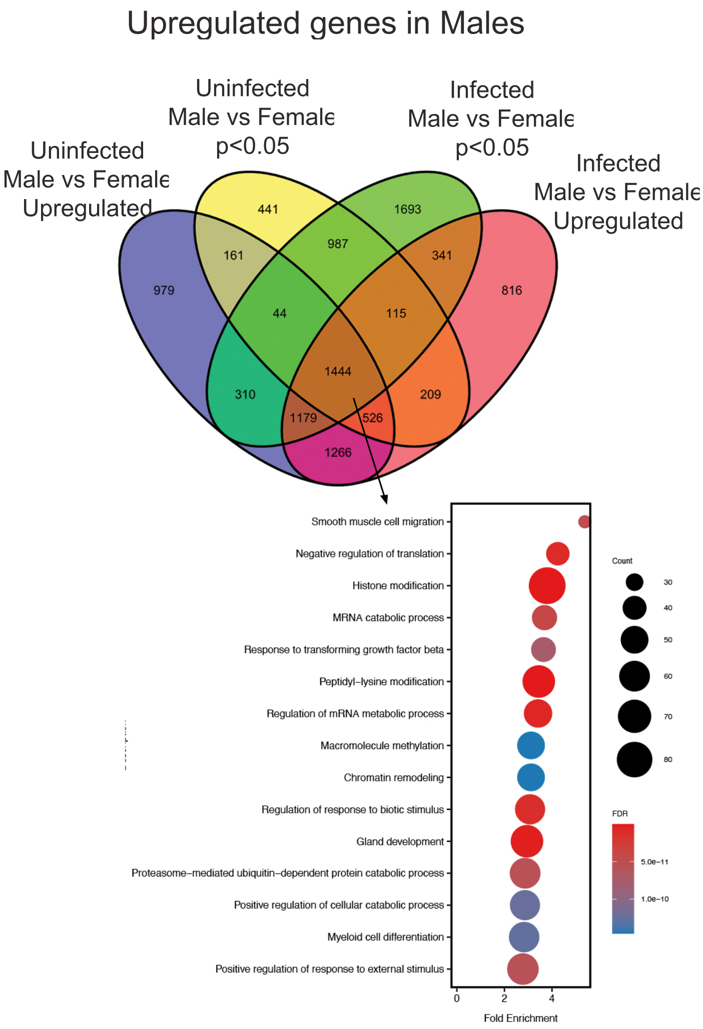


B

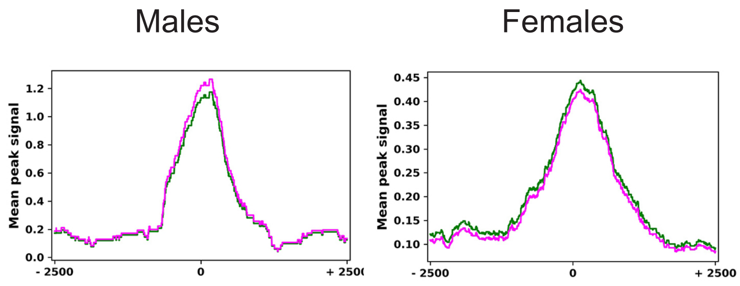


A) Differential expression analysis of genes upregulated in both uninfected and infected males, to generate the male-specific transcriptome adjusted p-value <0.05 along with pathway analysis of the 1444 genes significantly upregulated in males regardless of infection status. B) Plot of average of rlog normalized peaks counts (normalized by DESeq2) for all genes in each comparison that have significant peaks around their TSS. Green is uninfected magenta is infected.
